# Novel cytokinetic ring components drive negative feedback in cortical contractility

**DOI:** 10.1101/633743

**Authors:** Kathryn Rehain Bell, Michael E. Werner, Anusha Doshi, Daniel B. Cortes, Adam Sattler, Thanh Vuong-Brender, Michel Labouesse, Amy Shaub Maddox

## Abstract

Actomyosin cortical contractility drives many cell shape changes including cytokinetic furrowing. While positive regulation of contractility is well characterized, counterbalancing negative regulation and mechanical brakes are less well understood. The small GTPase RhoA is a central regulator, activating cortical actomyosin contractility during cytokinesis and other events. Here we report how two novel cytokinetic ring components, GCK-1 and CCM-3, participate in a negative feedback loop among RhoA and its cytoskeletal effectors to inhibit contractility. GCK-1 and CCM-3 are recruited by active RhoA and anillin to the cytokinetic ring, where they in turn limit RhoA activity and contractility. This is evidenced by increased RhoA activity, anillin and non-muscle myosin II in the cytokinetic ring, and faster cytokinetic furrowing, following depletion of GCK-1 or CCM-3. GCK-1 or CCM-3 depletion also reduced RGA-3 levels in pulses, and increased baseline RhoA activity and pulsed contractility during zygote polarization. Together, our findings suggest that GCK-1 and CCM-3 regulate cortical actomyosin contractility via negative feedback.

**Summary:** Novel cytokinetic ring proteins, the Ste20 family kinase GCK-1 and its heterodimeric cofactor Cerebral Cavernous Malformations-3, close a negative feedback loop involving the RhoA GAP RGA-3/4, RhoA, and its cytoskeletal effector anillin to limit actomyosin contractility in cytokinesis and during polarization of the *C. elegans* zygote.

## Introduction

Following faithful replication and segregation of the genome, nascent daughter nuclei are partitioned into individual daughter cells during cytokinesis (Green et al., 2012; Pollard and O’Shaughnessy, 2019). The spatiotemporal coupling of this partitioning with chromosome segregation is achieved by signaling from the anaphase spindle to elicit a zone of active RhoA GTPase at the equatorial cortex (Basant and Glotzer, 2018). RhoA activity initiates a cascade of downstream effects ultimately resulting in the polymerization of filamentous actin (F-actin) and activation of the motor protein non-muscle myosin II (NMM-II). F-actin and NMM-II form the structural basis of the cytokinetic ring along with many other components, including anillin (D’Avino et al., 2015). Anillin binds F-actin, NMM-II, RhoA and other structural and regulatory ring components, thus acting as a scaffold (D’Avino, 2009; Piekny and Maddox, 2010). Once the ring is assembled, it constricts, drawing the plasma membrane into a furrow that partitions the cytoplasm (Cheffings et al., 2016). Complete cytokinetic ring closure is essential for cells to maintain proper ploidy. Failure results in the formation of a polypoid cell that can undergo apoptosis or cancerous transformation (Lacroix and Maddox, 2012).

RhoA activity that promotes cytokinesis as well as contractility in many other biological processes is regulated by the delicate balance of activating GEFs and inhibiting GAPs as well as positive and negative feedback loops (Bement et al., 2015; Bischof et al., 2017; Goryachev et al., 2016; Michaux et al., 2018; Nishikawa et al., 2017). The main RhoA GAP during cytokinesis, RGA-3/4, plays a central role in negative feedback regulation of pulsed contractions during polarization of the *C. elegans* zygote, but the mechanism of negative feedback regulation during cytokinesis remains poorly understood (Mayer et al., 2010; Michaux et al., 2018; Naganathan et al., 2014; Naganathan et al., 2018; Schmutz et al., 2007; Schonegg et al., 2007; Zanin et al., 2013; Zhang and Glotzer, 2015).

To gain new insights into the mechanisms of cytokinetic ring constriction, we identified novel anillin-interacting proteins and found Germinal Center Kinase – 1 (GCK-1), a serine/threonine kinase related to budding yeast Ste20 (sterile-20). GCK-1 is the only *C. elegans* orthologue of the mammalian germinal center kinase III subfamily, which is implicated in apoptosis, proliferation, polarity and cell motility (Rehain-Bell et al., 2017; Schouest et al., 2009; Yin et al., 2012; Zheng et al., 2010). One well characterized interaction partner of GCK-1 is CCM-3, which is thought to recruit GCK-1 to the STRIPAK complex (Hwang and Pallas, 2014). Together these proteins are thought to regulate endothelial integrity in part by negatively regulating RhoA (Borikova et al., 2010; Richardson et al., 2013; Zheng et al., 2010). We and others characterized the roles of *C. elegans* GCK-1 and its co-factor Cerebral Cavernous Malformations – 3 (CCM-3; collectively, GCK-1/CCM-3) in maintaining structural integrity of the oogenic syncytial germline (Pal et al., 2017; Rehain-Bell et al., 2017). GCK-1/CCM-3 promote the stability of the intercellular bridges that connect developing oocytes with a shared cytoplasm by limiting the abundance of proteins that promote contractility (including anillin and NMM-II (NMY-2)) (Pal et al., 2017; Rehain-Bell et al., 2017). We proposed that GCK-1/CCM-3 suppress anillin (ANI-1) and NMY-2 localization by inhibiting RhoA, as their vertebrate homologs are known to do (Rehain-Bell et al., 2017; Richardson et al., 2013).

The identification of novel anillin-interacting proteins that limit contractility fit logically with their localization to stable intercellular bridges. However, we and others noted that GCK-1/CCM-3 also enrich on the dynamic, contractile cytokinetic ring in the *C. elegans* zygote (Pal et al., 2017; Rehain-Bell et al., 2017). Here, we report investigating the regulation of contractility in the *C. elegans* zygote by GCK-1/CCM-3. We found that on the contractile ring, GCK-1/CCM-3 limit the abundance of “contractility” proteins. Partial depletion of GCK-1/CCM-3 boosts contractility during cytokinesis and polarization of the zygote. GCK-1/CCM-3 localize to the cytokinetic ring and cortical pulses downstream of the master regulator RhoA (RHO-1 in *C. elegans*) and of anillin (ANI-1), and also influence the abundance of active RhoA, likely through promoting the cortical recruitment or retention of RGA-3. Therefore, we conclude that GCK-1/CCM-3 are novel components of negative feedback in the cytokinetic ring. These findings advance the growing body of work showing that contractile networks in cells are not only activated by positive regulation, but also contain structural “brakes” and regulatory time-delayed negative feedback important for turnover and dynamics (Bement et al., 2015; Bischof et al., 2017; Dorn et al., 2016; Goryachev et al., 2016; Khaliullin et al., 2018; Michaux et al., 2018; Nishikawa et al., 2017).

## Results

### GCK-1 and CCM-3 regulate each other’s stability and cortical targeting

*C. elegans* GCK-1 and CCM-3 are known to interact, and their mammalian homologs form a heterodimer (Ceccarelli et al., 2011; Lant et al., 2015; Xu et al., 2013; Zhang et al., 2013). The domains that allow for heterodimerization of GCK III subfamily members and CCM3 are conserved in *C. elegans* GCK-1 and CCM-3 indicating that these proteins heterodimerize as well (Ceccarelli et al., 2011). We first tested the dynamics of GCK-1/CCM-3 localization and their interdependence during cytokinesis. We followed the localization of fluorescently-tagged GCK-1 and CCM-3 expressed under the control of their own promoters (Pal et al., 2017; Rehain-Bell et al., 2017). Both are present in the cytoplasm and enrich in the cytokinetic ring during anaphase (Figure 1A and B). We then depleted GCK-1 or CCM-3 by RNAi to test the requirement of each for localization of the other to the cytokinetic ring. Levels of GCK-1 or CCM-3 in the cytokinetic ring were significantly reduced following depletion of CCM-3 or GCK-1, respectively (Figure 1A, A’, B, B’). These results demonstrate that GCK-1 and CCM-3 are interdependent for their enrichment on the cytokinetic ring and support the idea that they act as a complex during cytokinesis.

**Figure 1.**
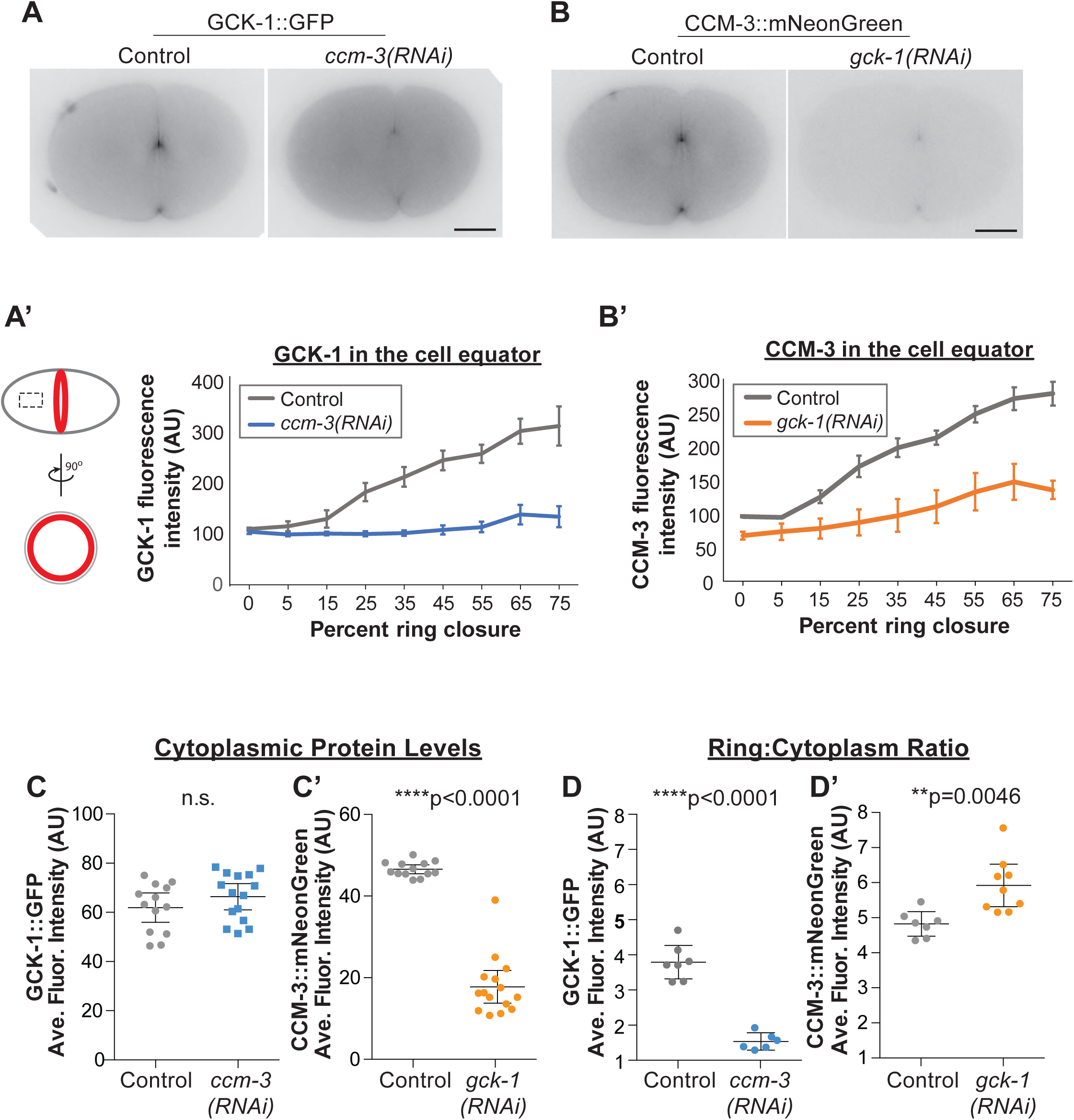
Mutual regulation of GCK-1 and CCM-3 enrichment on the cytokinetic ring. A) Representative single plane images of control and *ccm-3(RNAi)* embryos expressing GCK-1::GFP. A) Representative single plane images of control and *gck-1(RNAi)* embryos expressing CCM-3::mNeonGreen. A’ and B’) Quantitation of average fluorescence intensity per micron relative to the perimeter of the cytokinetic ring in control (n=7 embryos) and depleted (n= 9 or 6 embryos (CCM3 or GCK-1 depleted, respectively)) conditions. C and C’) Cytoplasmic levels of GCK-1::GFP (C) and CCM-3::mNeonGreen (C’) in control and depleted cells. D and D’) Enrichment of GCK-1::GFP (D) and CCM-3::mNeonGreen (D’) on the cytokinetic ring relative to the cytoplasm in control and depleted cells. Scale bars are 10 μm in all images in all figures. Error bars are SE in all figures unless otherwise indicated.

We next tested whether GCK-1 and CCM-3 affect each other’s localization to the cytokinetic furrow via active recruitment or effect on protein level. To do so, we assessed the abundance of cytoplasmic GCK-1 or CCM-3 following depletion of the other (Figure 1C and C’). CCM-3 depletion did not significantly affect the cytoplasmic levels of GCK-1::GFP (Figure 1C). Thus, we concluded that CCM-3 is not required for GCK-1 protein stability. Depletion of GCK-1, however, significantly decreased cytoplasmic levels of CCM-3::mNeonGreen (Figure 1C’) suggesting that GCK-1 is required for CCM-3 stability. Consistently, we found that cytokinetic ring enrichment, the ratio of cytokinetic ring and cytoplasmic protein levels, of CCM-3::mNeonGreen increased slightly (∼20%) following GCK-1 depletion compared to control, indicating approximately equal reduction of both the ring and cytoplasmic protein pools and little effect on ring targeting (Figure 1D’). Conversely, GCK-1::GFP enrichment on the cytokinetic ring relative to the cytoplasm was significantly decreased (∼60% decrease) following CCM-3 depletion (Figure 1D). Together these results suggest that CCM-3 targets GCK-1 to the cytokinetic ring, while GCK-1 promotes CCM-3 ring localization at least partly by regulating CCM-3 protein level. *GCK-1 and CCM-3 localize downstream of RhoA*

RhoA is the master regulator necessary for recruitment and activation of cytokinetic ring components in animal cells (Basant and Glotzer, 2018; Jordan and Canman, 2012; Piekny et al., 2005). To test whether GCK-1/CCM-3 localize downstream of RhoA (RHO-1 in *C. elegans*), we performed time-lapse imaging of GCK-1::GFP and CCM-3::mNeonGreen on the cell cortex during anaphase. Shortly following anaphase onset in control cells, both GCK-1 and CCM-3 localize to large cortical foci in the anterior of the embryo and the cell equator (Figure 2A and Supplemental Figure 1A. This localization pattern mirrors that of many known cytokinetic ring components such as active RHO-1, anillin (ANI-1) and NMY-2 (Maddox et al., 2005; Maddox et al., 2007; Motegi et al., 2006; Schonegg et al., 2007; Velarde et al., 2007; Werner et al., 2007). In fact, GCK-1 and CCM-3 colocalized with NMY-2 during anaphase (Figure 2A, Supplemental movie 1 and Supplemental Figure 1A). When the levels of active RHO-1 were reduced by partially depleting its main activator during cytokinesis the RhoGEF ECT-2, the levels of cortical GCK-1 or CCM-3 at the cell equator during anaphase were significantly reduced (Figure 2B and C, Supplemental movie 2, and Supplemental Figure 1B and C). We conclude that GCK-1/CCM-3 depend on active RhoA for recruitment to the cytokinetic ring.

**Figure 2.**
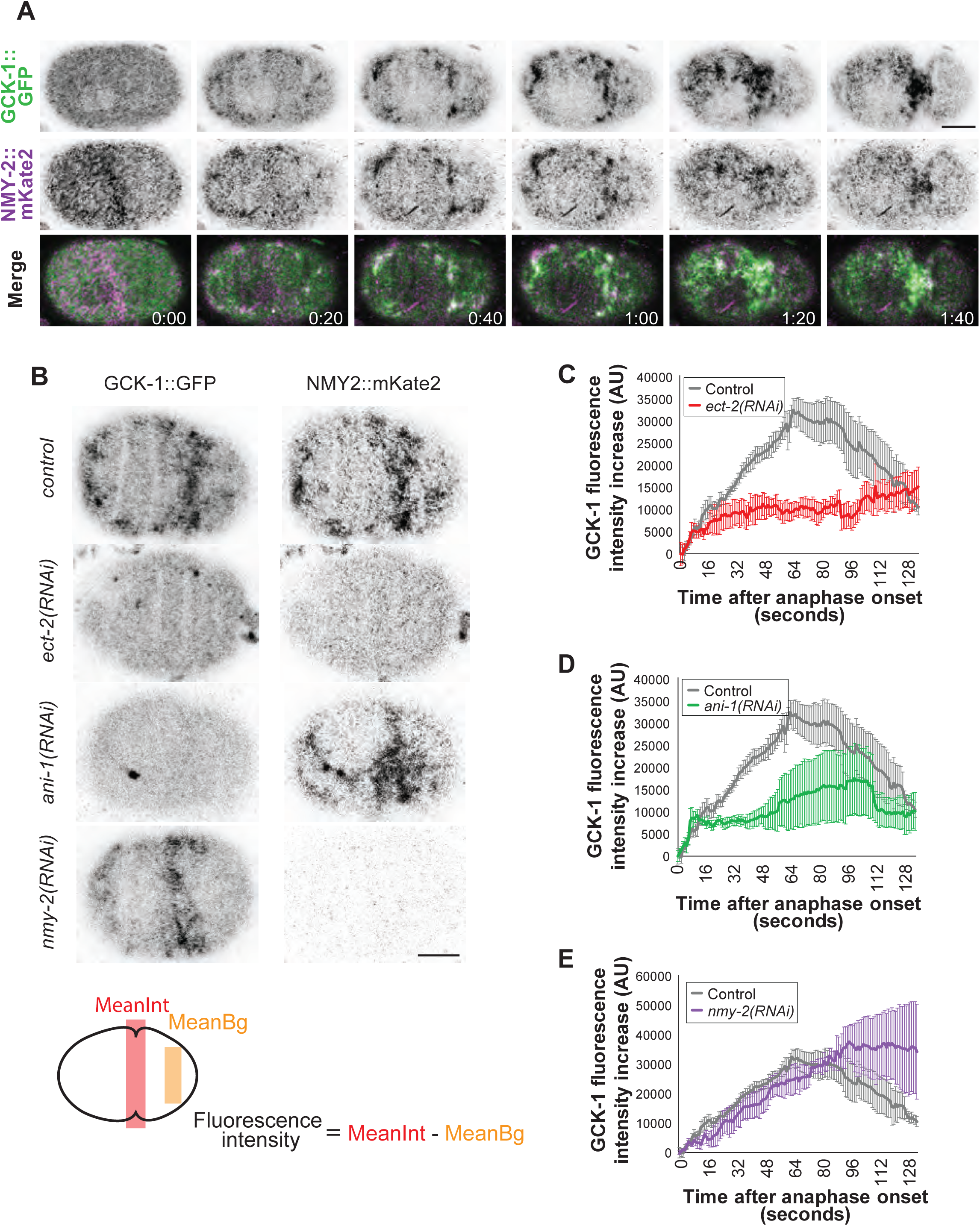
GCK-1 localization is dependent on active RHO-1 and ANI-1 but not NMY-2. A) Representative images of GCK-1::GFP and NMY-2::mKate localization during anaphase. B) Representative images of embryos expressing both GCK-1::GFP and NMY-2::mKate in control, *ect-2(RNAi), ani-1(RNAi)* and *nmy-2(RNAi)* embryos 60 seconds after anaphase onset. C-E) Background-adjusted equatorial GCK-1::GFP levels normalized to that at anaphase onset in control, *ect-2(RNAi)* (C), *ani-1(RNAi)* (D) and *nmy-2(RNAi)* (E) embryos (n=6 for all conditions).

RhoA effector proteins including Diaphanous-family formins, Rho-kinase, and anillin bind RhoA-GTP directly and independently, whereas downstream factors such as F-actin, NMY-2, and septins depend on these effectors (Heasman and Ridley, 2008; Piekny et al., 2005; Piekny and Maddox, 2010). To determine if GCK-1 localization is dependent on other cytokinetic ring components, we depleted ANI-1 or NMY-2 by RNAi and assessed GCK-1 and CCM-3 levels in the cytokinetic ring. Depletion of ANI-1, but not NMY-2, caused a significant reduction in GCK-1 and CCM-3 on the cell equator (Figure 2B, D and E, Supplemental movies 3 and 4 and Supplemental Figure 1B, D and E). Taken together, these results demonstrate that GCK-1 and CCM-3 are recruited to the cytokinetic ring downstream of active RHO-1 and ANI-1 but not NMY-2.

### GCK-1 and CCM-3 limit contractility in cytokinesis

We next tested whether, as in the stable intercellular bridges of the *C. elegans* germline, GCK-1 or CCM-3 limit contractility in the highly contractile zygote. We began by assessing embryonic viability, and found that depletion of GCK-1 or CCM-3 leads to 11% or 13% embryonic lethality, respectively (Figure 3A). To explore if this embryonic lethality could be due to defects in cytokinesis in early (< 100 cells) embryos, we strongly depleted GCK-1 or CCM-3 and quantified the prevalence of multinucleation. A similar percentage of GCK-1 (15%) or CCM-3 (20%) depleted embryos contained at least one multinucleated blastomere (Figure 3B). However, cytokinesis failure was rare; only about 0.5% of blastomeres were multinucleate (Figure 3C). As reported previously, strong depletion of GCK-1 or CCM-3 caused severe defects in germline organization leading to smaller and fewer embryos (Green et al., 2011; Pal et al., 2017; Priti et al., 2018; Rehain-Bell et al., 2017; Schouest et al., 2009). As such, embryos assessed for cytokinesis defects were significantly smaller (Supplemental Figure 2A) and a correlation between embryo size and frequency of multinucleation suggested that the latter could at least partially be a secondary effect of abnormal embryo size (Supplemental Figure 2B).

**Figure 3.**
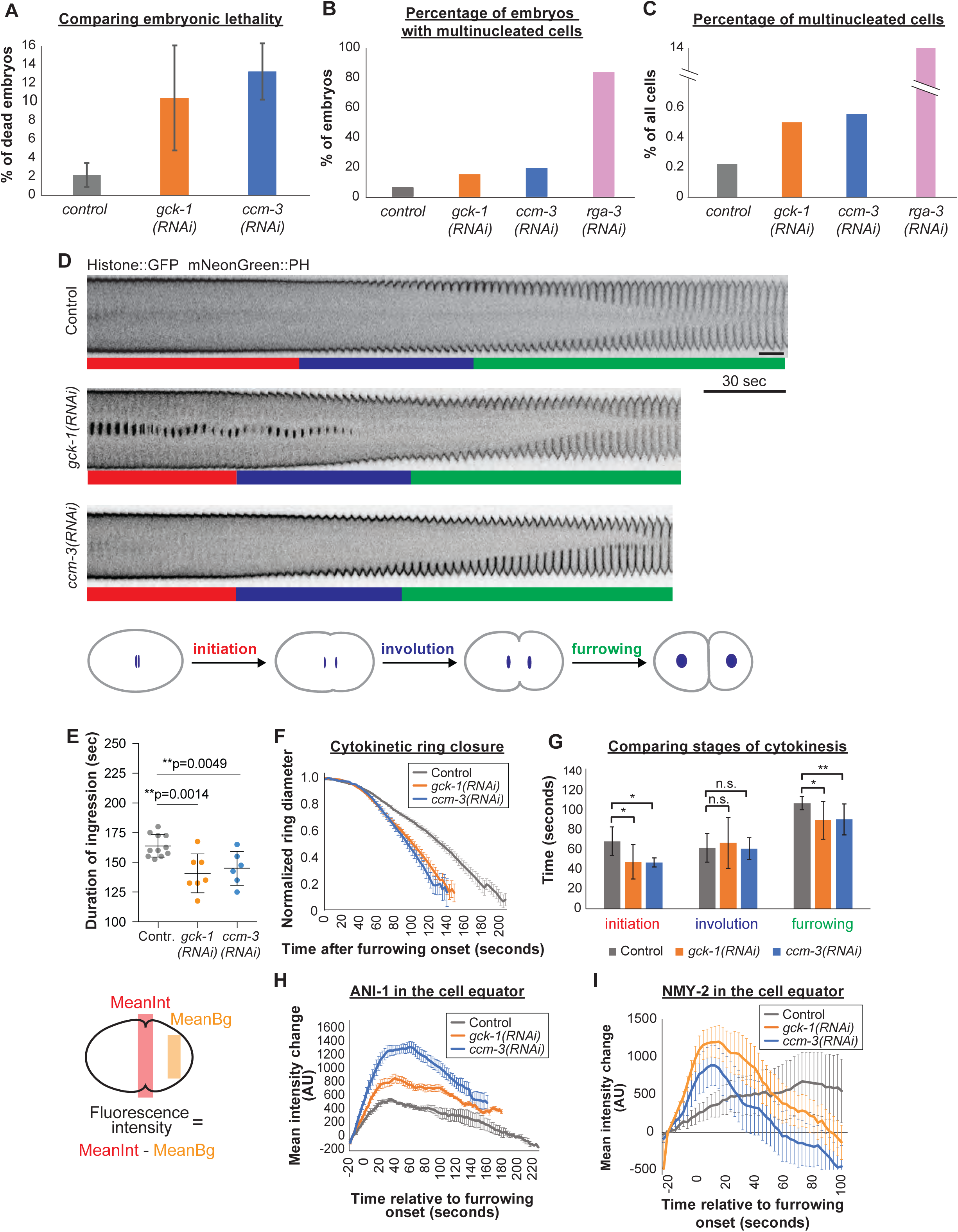
Depletion of GCK-1 or CCM-3 causes increased cytokinetic ingression speed and increase the amount of NMY-2 and ANI-1 in the furrow. (A) Quantification of embryonic lethality in control embryos or after depletion of *gck-1(RNAi)* and *ccm-3(RNAi)*. (B) Percent of embryos with at least one multinucleated cell in control embryos or after strong gck-1(RNAi), *ccm-3(RNAi)* or *rga-3(RNAi)* (n>80 embryos for all conditions). (C) Quantification of the amount of multinucleated cells as a percentage of total cells in embryos quantified in B. (D) Representative montage of HIS::GFP and mNeonGreen::PH in the furrow region over time in control, *gck-1(RNAi)* and *ccm-3(RNAi)* embryos. Colored bars represent the timing of distinct phases of cytokinesis shown in the schematic below. Initiation: anaphase onset - onset of furrow ingression. Involution: onset of ingression - appearance of a doubled membrane. Furrowing: the appearance of doubled membrane - complete ring closure. E) Quantification of the duration of interval between furrowing onset and ring closure. F-G) Cytokinetic ring closure dynamics in control (n=9), *gck-1(RNAi)* (n=7) and *ccm-3(RNAi)* (n=6) embryos measured by following closure dynamics (F) or subdividing cytokinesis timing into 3 distinct stages (G) using a strain expressing mNeonGreen::ANI-1 and NMY-2::mKate2. H) Quantification of background-adjusted equatorial mNeonGreen::ANI-1 levels relative to equatorial levels 20 seconds prior to onset of furrow ingression in control (n=6), *gck-1(RNAi)* (n=9) or *ccm-3(RNAi)* (n=9) embryos. I) Quantification of background-adjusted equatorial NMY-2::GFP levels relative to equatorial levels 20 seconds prior to onset of furrow ingression in control (n=9), *gck-1(RNAi)* (n=10) and *ccm-3(RNAi)* (n=12) embryos. Error bars are SD in A, E and G.

To study the role of GCK-1/CCM3 in cytokinesis while avoiding the confounding factor of abnormal embryo size, we performed feeding RNAi for 24 hours to partially (>83%) deplete embryos of CCM-3 or GCK-1, which maintained normal embryo size (Supplemental Figure 2C). Under these conditions, cytokinetic furrows ingressed to completion in all zygotes observed, and furrowing completed significantly faster than controls, largely due to increased maximal ingression speed of the cytokinetic ring (Figure 3D-G). This suggests that GCK1/CCM-3 attenuate contractility during cytokinetic ring constriction.

To determine how GCK-1/CCM-3 affect cytokinetic ring ingression kinetics, we examined how they affect recruitment of cytokinetic ring components to the cell equator. The levels of both NMY-2 and ANI-1 at the cell equator were significantly increased following GCK-1 or CCM-3 depletion (Figure 3H and I). This result suggests that GCK-1 and CCM-3 attenuate ingression speed by limiting the amount of proteins that drive contractility in the cytokinetic ring.

Taken together, these results demonstrate that GCK-1/CCM-3 limit the amount of ANI-1 and NMY-2 at the cell equator and negatively regulate the kinetics of cytokinetic ring ingression.

### GCK-1 and CCM-3 negatively regulate cortical contractility during polarization

Prior to the first mitotic division of the *C. elegans* zygote, a highly contractile actomyosin network spans the embryonic cortex and becomes progressively polarized to the embryo anterior. To test whether GCK-1/CCM-3 limit zygote contractility not only in cytokinesis but also more generally, we next examined the role of GCK-1/CCM-3 during polarization, when a transient feature called the pseudocleavage furrow ingresses at the boundary of the highly contractile anterior and the more compliant posterior cortex, and regresses. The positioning and regulation of the pseudocleavage furrow are distinct and independent from those of the cytokinetic furrow that ingresses about 10 minutes later (Cowan and Hyman, 2007; Werner and Glotzer, 2008) (Figure 4A). Consistent with a potential role before cytokinesis, we observed that GCK-1/CCM-3 enrich in cortical patches during polarization, co-localizing with other contractile components such as NMY-2 and ANI-1 (Figure 4B, Supplemental movie 5 and Supplemental Figure 3A and B). Similar to our observations during anaphase this localization is dependent on ECT-2 and ANI-1 but not NMY-2 (Supplemental Figure 3C and D).

**Figure 4.**
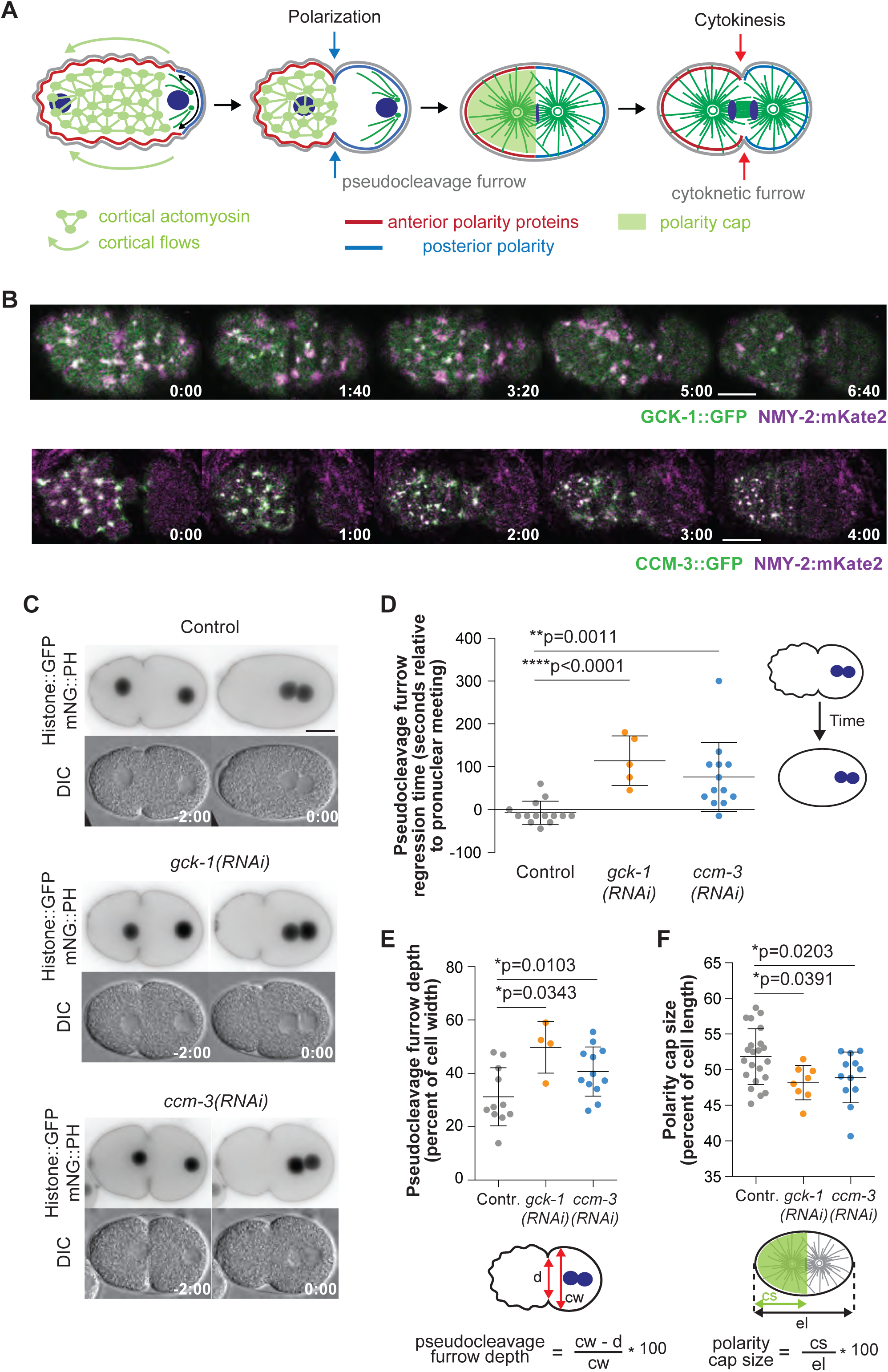
GCK-1 localizes to cortical patches during polarization and inhibits cortical contractility. A) Schematic representation cellular morphology and distribution of relevant protein complexes in the *C. elegans* zygote. B) Representative images of single plane cortical imaging of embryos expressing GCK-1::GFP and NMY-2::mKate2, or CCM-3::mNeonGreen and NMY-2::mKate2. C) Representative fluorescent images of control, *gck-1(RNAi) and ccm-3(RNAi)* embryos expressing HIS::GFP and mNeonGreen::PH (Top) and corresponding DIC images (bottom). Left: maximal pseudocleavage furrow ingression depth; Right: at pronuclear meeting. D) Quantification of the time interval between pronuclear meeting and complete regression of the pseudocleavage furrow in control, *gck-1(RNAi)* and *ccm-3(RNAi)* embryos. (E) Quantification of maximal pseudocleavage ingression as a percentage of cell width in control, *gck-1(RNAi)* and *ccm-3(RNAi)* embryos. (F) Quantification of the size of the anterior PAR domain at metaphase in control, *gck-1(RNAi)* and *ccm-3(RNAi)* embryos expressing PAR-6::mKate2. Error bars are SD.

To test whether GCK-1/CCM-3 affect contractility during polarization, we depleted either CCM-3 or GCK-1 by RNAi and measured the regression timing and depth of the pseudocleavage furrow, both of which reflect contractility during polarization (Reymann et al., 2016; Schmutz et al., 2007; Schonegg et al., 2007; Tse et al., 2012). The persistence of the pseudocleavage furrow was significantly increased in zygotes depleted of either CCM-3 or GCK-1 (Figure 4C and D and Supplemental movie 5). To control for possible defects in cell cycle progression, we also measured the time from pronuclear meeting to anaphase onset and found no statistically significant difference among control and embryos depleted of GCK-1 or CCM-3 indicating that cell cycle timing is not affected (Supplemental Figure 4A). Similarly, the maximal depth of pseudocleavage furrow as a percentage of embryo width was significantly increased following GCK-1 or CCM-3 depletion, when compared to controls (Figure 4C, E). An additional measure of cortical contractility during polarization is the size of the anterior polarity domain (Schonegg et al., 2007; Tse et al., 2012). The size of the anterior domain, defined by localization of PAR-6, was significantly reduced in embryos depleted of GCK-1 or CCM-3 when compared to controls (Figure 4F). This was not due to defects in polarity establishment or maintenance following depletion of CCM-3 or GCK-1, since asymmetric spindle positioning or the extent of asymmetric enrichment of anterior PAR proteins were not affected (Supplemental Figure 4B-B’’). The dependence of the anterior enrichment of GCK-1 and CCM-3 on PARs (Supplemental Figure 4C-G’’) further demonstrated that GCK-1 and CCM-3 are downstream of PAR-driven polarity. Taken together, these findings support the conclusion that GCK-1 and CCM-3 limit cortical contractility during polarization, in addition to in cytokinesis and germline intercellular bridges.

### GCK-1/CCM-3 limit RhoA activity in the cytokinetic ring and during pulsed contractility

GCK-1/CCM-3 homologues in cultured human endothelial cells and mouse neocortex negatively regulate RhoA (Borikova et al., 2010; Louvi et al., 2014; Zheng et al., 2010), but we showed that they are recruited to the cytokinetic ring downstream of RHO-1 activity and ANI-1. We reasoned that this series of recruitment interdependencies and protein activities could constitute a negative feedback loop in which RHO-1 recruits ANI-1, which then recruits GCK-1/CCM-3, which in turn limits RHO-1 activity and contractility. To test this hypothesis, we first examined the pulsed cortical contractions that occur during embryo polarization. These cycles of quiescence, activation, contraction, and finally relaxation and disassembly are a powerful model to study feedback loops in the regulation of contractility (Michaux et al., 2018; Naganathan et al., 2018; Nishikawa et al., 2017; Reymann et al., 2016). RHO-1 activity in *C. elegans* has been assessed using a biosensor composed of the C-terminal third of ANI-1 which contains, among several functional elements, its RHO-1 binding domain (Sun et al., 2015). Since GCK-1/CCM-3 and ANI-1 reciprocally affect the other’s localization and/or stability (Figure 2B and D and Figure 3H), we avoided the use of the ANI-1 based biosensor, and generated a fluorescently-tagged Rho-kinase (LET-502) expressed from its endogenous locus to employ as a biosensor for RHO-1 activity. Rho-kinase is a well-characterized RhoA effector that preferentially associated with the activated form of RhoA (Matsui et al., 1996). To characterize this probe, we increased or decreased RHO-1 activity by depleting its inactivator RhoGAP RGA-3/4 or activator RhoGEF ECT-2, respectively. As expected, recruitment of LET-502 to the cell equator increased following RGA-3/4 depletion and decreased following ECT-2 depletion (Figure 5A-C and Supplemental movies 6-8).

**Figure 5.**
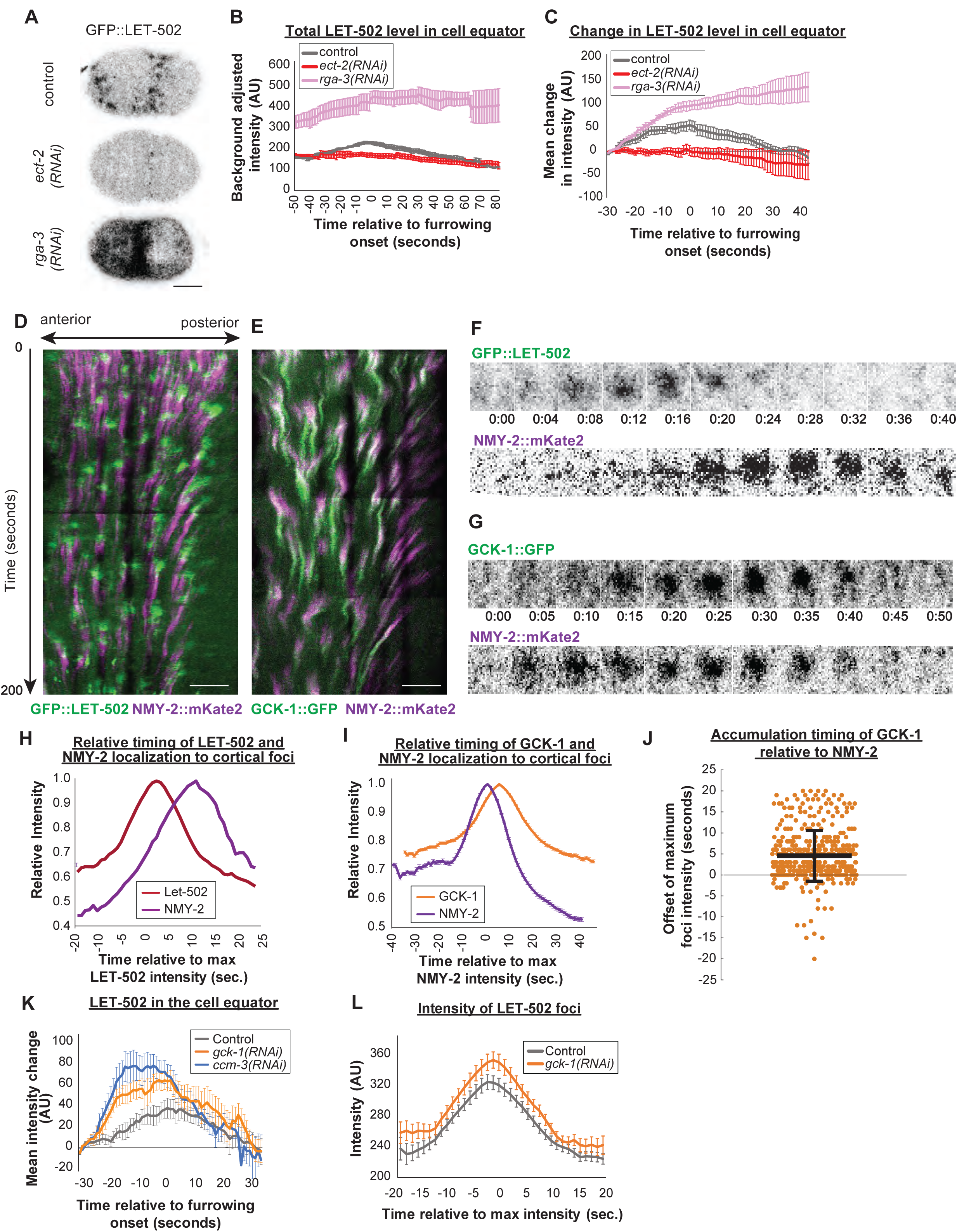
GCK-1 localizes to cortical foci after NMY-2 and limits RHO-1 activity on the cortex. (A) Representative images of endogenous GFP::LET-502 at onset of ingression in control, *ect-2(RNAi)* or *rga-3/4(RNAi)* embryos. B) Background-adjusted total equatorial GFP::LET-502 fluorescence levels in control embryos (n=7) and embryos with increased (*rga-3/4(RNAi)* (n=6)) or decreased (*ect-2(RNAi)* (n=6)) active RHO-1 levels. C) Relative increase in background adjusted total equatorial GFP::LET-502 fluorescence levels in control embryos and embryos with increased (*rga-3/4(RNAi)*) or decreased (*ect-2(RNAi)*) active RHO-1 levels. D) Kymograph showing pulsed GFP::LET-502 and NMY-2::mRFP localization during embryo polarization. E) Kymograph showing pulsed GCK-1::GFP and NMY-2::mKate2 localization during embryo polarization. F) Magnified view of GFP::LET-502 and NMY-2::mRFP localization to a single focus. G) Magnified view of GCK-1::GFP and NMY-2::mKate2 localization to a single focus. H) Mean accumulation profile of GFP::LET-502 and NMY-2:mRFP in cortical patches I) Mean accumulation profile of GCK-1::GFP and NMY-2:mKate2 in cortical patches. J) Time delay in maximal GCK-1::GFP accumulation relative to NMY-2::mKate2 localization. Data points = individual foci; n=236 total foci from a total of 5 embryos for H and n=423 total foci from a total 8 embryos for I and J. K) Quantification of background-adjusted equatorial GFP::LET-502 levels relative to equatorial levels 30 seconds prior to onset of furrow ingression in control (n=7), *gck-1(RNAi)* (n=10) and *ccm-3(RNAi)* (n=10) embryos. L) Mean fluorescence intensity profile of GFP::LET-502 in cortical patches from control and *gck-1(RNAi)* embryos. Error bars are SE except for panel J, where error bars are SD.

We next tested whether GCK-1/CCM-3 localization dynamics are consistent with a role in negative feedback during pulsed contractility. Consistent with previous observations, local enrichment of active RHO-1, as visualized via our LET-502 biosensor, precedes the localization of NMY-2, ANI-1 and RGA-3, all of which localize concurrently (Michaux et al., 2018) (Figure 5D, F and H, Supplemental movie 9). We imaged cells co-expressing GFP-tagged GCK-1 and RFP-tagged NMY-2 with high temporal resolution and tracked cortical patches during polarization to determine the kinetics of GCK-1 localization relative to the accumulation of NMY-2. The maximal level of GCK-1 in cortical patches occurred approximately 3.5 seconds after that of NMY-2 and thus about 13 seconds after active RHO-1 recruitment, consistent with a role in time-delayed negative feedback regulation of RHO-1 and contractility (Figure 5E, G, I and J and Supplemental movie 10).

We predicted that if GCK1/CCM3 feed back negatively on RHO-1, depletion of GCK-1/CCM-3 would increase levels of active RHO-1. This prediction was substantiated by the positive control that depletion of RGA-3/4 increased the baseline RHO-1 activity level during pulsed contractility (Figure 5L and 6D). We tested the effects of depleting GCK-1/CCM-3 on the abundance of active RHO-1, in the cell equator and in cortical pulses during polarization. The abundance of LET-502 was significantly increased following GCK-1 or CCM-3 depletion, when compared to controls, in both the cell equator and cortical pulses (Figure 5K and L). Together, our results support the conclusion that GCK-1/CCM3 are novel cytokinetic ring components recruited downstream of RHO-1 (via ANI-1) that in turn feed back negatively on RhoA activity (Figure 6G).

**Figure 6.**
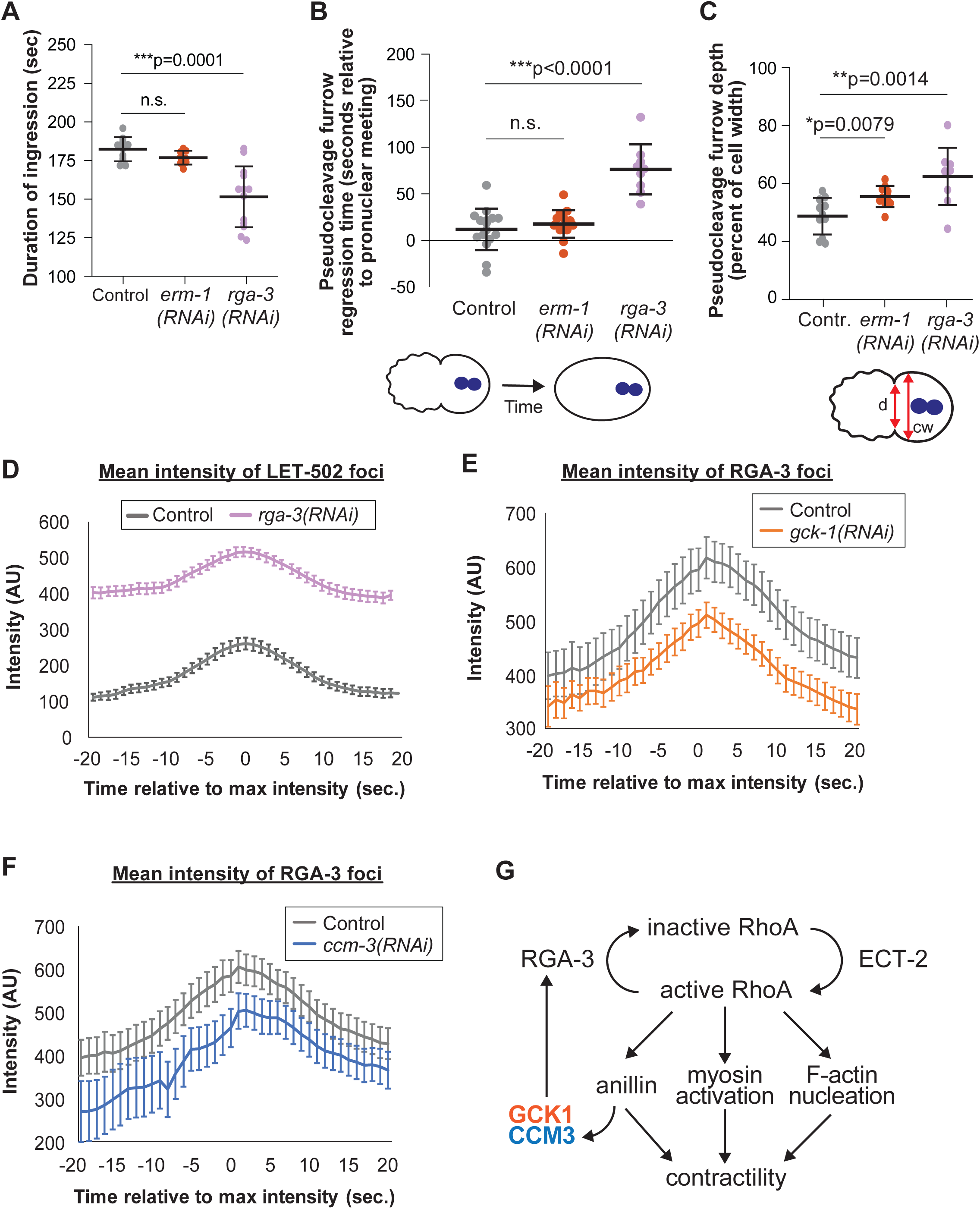
GCK-1/CCM-3 contribute to negative feedback regulation of RHO-1 by promoting RGA-2 localization. A) Quantification of the duration of interval between furrowing onset and ring closure in control, *erm-1(RNAi)* or weak *rga-3(RNAi)* embryos. B) Time interval between pronuclear meeting and complete regression of the pseudocleavage furrow in control, *erm-1(RNAi)* or weak *rga-3(RNAi)* embryos. (C) Quantification of maximal pseudocleavage ingression as a percentage of cell width in control, *erm-1(RNAi)* or weak *rga-3(RNAi)* embryos. (D) Mean fluorescence intensity profile of GFP::LET-502 in cortical patches from control and weak *rga-3(RNAi)* embryos (n=9 embryos each for a total of 277 and 304 patches respectively). (E) Mean fluorescence intensity profile of RGA-3::GFP in cortical patches from control (n=9 embryo; 241 patches total) and *gck-1(RNAi)* embryos (n=8 embryos; 155 patches total). (F) Mean fluorescence intensity profile of RGA-3::GFP in cortical patches from control and *ccm-3 (RNAi)* embryos (n=8 embryos 140 total patches). (G) Summary diagram of relationship of GCK-1/CCM-3 to canonical cytokinetic contractility regulation network. Error bars are SD in A-C; all others are SE.

### GCK-1/CCM-3 regulate RhoA activity by promoting RGA-3/4 recruitment

What is the molecular mechanism of RhoA inhibition downstream of GCK-1/CCM-3? In zebrafish and human cultured cells, homologues of GCK-1 phosphorylate moesin, an ezrin-radixin-moesin (ERM) family protein, leading to a reduction of RhoA activity (Zheng et al., 2010). To determine whether GCK-1 regulates RHO-1 activity by a similar mechanism in the *C. elegans* zygote, we tested whether depletion of the only *C. elegans* ERM protein, ERM-1, affects cytokinetic furrow ingression kinetics or contractility during polarization (Gobel et al., 2004; Van Furden et al., 2004). Neither cytokinetic furrow kinetics nor pseudocleavage regression timing was significantly different between control and ERM-1 depleted cells, and the pseudocleavage furrow was only slightly deeper following ERM-1 depletion, suggesting that ERM-1 is not the main target of GCK-1/CCM-3 during polarization or cytokinesis (Figure 6 A-C).

Another potential mediator of RHO-1 inhibition by GCK-1/CCM-3 is the RhoGAP RGA-3/4, a known negative regulator of RhoA activity during both polarization and cytokinesis that mediates negative feedback regulation during pulsed contractions (Michaux et al., 2018; Schmutz et al., 2007; Schonegg et al., 2007; Zanin et al., 2013). Severe reduction or elimination of RGA-3/4 levels following thorough RNAi or in homozygous *rga-3Δ, rga-4Δ* double mutants leads to ectopic contractility, cytokinetic furrow positioning defects, and undetectable active RHO-1 pulses, confounding quantitative comparisons with the contractile phenotypes following GCK-1/CCM-3 depletion (Michaux et al., 2018; Schmutz et al., 2007; Schonegg et al., 2007; Zanin et al., 2013). Interestingly, in heterozygous *rga-3/4 (+/Δ, +/Δ)* double mutants cortical contractility still increases but pulses of active RHO-1 persist (Michaux et al., 2018). We used partial RGA-3/4 depletion to quantify RHO-1 pulses when RHO-1 activity is increased. We observed an increase in baseline active RhoA levels in cortical pulses without a significant effect on the length of the pulse (Figure 6D). Hence, weak RGA-3/4 depletion enhanced contractility during both cytokinesis and polarization in ways that closely resembled the defects observed following GCK-1/CCM-3 depletion (Figure 3E, Figure 4D and E and Figure 6A-C).

We next tested whether GCK-1/CCM-3 depletion affected cortical abundance of RGA-3/4. Indeed, depletion of GCK-1 or CCM-3 significantly reduced GFP::RGA-3 levels in cortical pulses (Figure 6 E and F). These results support the idea that GCK-1/CCM-3 regulate RhoA activity by promoting the cortical recruitment of the RhoGAP RGA-3. Together, our results define a negative feedback loop among RHO-1, ANI-1, GCK-1/CCM-3 and RGA-3/4 (Figure 6G).

## Discussion

We characterized two novel cytokinetic ring proteins and their roles in regulating actomyosin contractility during cytokinesis. We found that GCK-1/CCM-3 are recruited downstream of active RHO-1 and the scaffold protein ANI-1, suggesting they are indirect RHO-1 effectors. However, GCK-1/CCM-3 also limit RHO-1 activity and actomyosin contractility during pulsed contractility, polarization, and cytokinesis, in agreement with what we and others showed for the stable actomyosin cortex of the *C. elegans* syncytial germline (Pal et al., 2017; Priti et al., 2018; Rehain-Bell et al., 2017). We report a potential mechanism by which GCK-1/CCM-3 inhibit Rho: they help recruit or retain the RhoGAP RGA-3/4 at the cortex. Therefore, we propose that GCK-1/CCM-3 contribute to a negative feedback loop acting on the RHO-1-regulated actomyosin cytoskeleton.

Our conclusion that GCK-1/CCM-3 contribute to negative feedback on RHO-1 activity is further supported by the delayed timing of GCK-1/CCM-3 localization to pulsed contractile cortical patches, with respect to active RHO-1 and NMY-2 and, by extension ANI-1, recruitment. Contractile pulses are driven by oscillations of RHO-1 activity, which drives F-actin assembly, which in turn promotes accumulation of the RhoGAP RGA-3/4 (Michaux et al., 2018; Nishikawa et al., 2017), which feeds back to inactivate RHO-1. To test how GCK-1/CCM-3 contribute to pulse dynamics, we first measured how pulses respond to a reduction of RHO-1 activity, since reduction and elimination of RGA-3/4 have qualitatively distinct effects (Michaux et al., 2018). To approximate the reduction in RGA-3/4 we observed following GCK-1/CCM-3 depletion, we examined pulses following depletion of RGA-3/4. In such cells, pulse dynamics were indistinguishable from those of control cells, but baseline Rho activity level was higher following RGA-3/4 depletion (Figure 6D). Similarly, depletion of GCK-1/CCM-3 increased the amount of baseline RHO-1 activity in cortical patches, at least partly by limiting the initial size of cortical patches, but did not change pulse dynamics (Figure 5L and Supplemental Figure 5). Higher baseline RhoA activity is expected to push an excitable system towards stable RhoA activation (Goryachev et al., 2016). We therefore predict that thorough loss of GCK-1/CCM-3, especially in combination with other RHO-1 activity gain-of-function perturbations, could cause constitutive cortical contractility and reduce the responsiveness of the cortex to spatial cues from the spindle.

Together, these observations suggest that GCK-1 and CCM-3 contribute to negative regulation of the cortical actomyosin cytoskeleton. This conclusion supports a growing consensus about negative feedback in pulsed contractility (Bement et al., 2015; Bischof et al., 2017; Dorn et al., 2016; Goryachev et al., 2016; Khaliullin et al., 2018; Michaux et al., 2018; Nishikawa et al., 2017). Interestingly though, it has not to our knowledge been demonstrated that negative feedback on RhoA contributes to normal cytokinetic dynamics. Our observations that negative feedback built into the cytokinetic ring limit its speed may reflect an advantage of the above-mentioned responsiveness of the system.

RHO-1 activity is required for equatorial enrichment of GCK-1/CCM-3. Several observations suggest that this is mediated by ANI-1: depletion of ANI-1 prevents GCK-1/CCM-3 equatorial enrichment (Figure 2B and D), ANI-1/anillin recruitment is downstream of active RHO-1/RhoA (Hickson and O’Farrell, 2008; Piekny and Glotzer, 2008; Piekny and Maddox, 2010; Straight et al., 2003), and GCK-1 co-immunoprecipitates with ANI-1 (Rehain-Bell et al., 2017). Anillins are large multi-domain scaffold proteins with many binding partners and putative roles in cytokinesis (Piekny and Maddox, 2010). It is possible that the increase in furrowing speed observed following partial ANI-1 depletion (Descovich et al., 2018) is due in part to decreased GCK-1/CCM-3 recruitment.

Thorough depletion of GCK-1/CCM-3 leads to severe germline defects and precludes the formation of normal embryos (Green et al., 2011; Pal et al., 2017; Rehain-Bell et al., 2017) (Priti et al., 2018; Schouest et al., 2009). The partial nature of our GCK-1/CCM-3 depletions likely explains the partial, but significant, effect on RGA-3/4 levels we observed (Figure 6E, F). GCK-1/CCM-3 have been anecdotally implicated in zygote cytokinesis and polarity (Pal et al., 2017). We quantitatively examined these events following various degrees of GCK-1/CCM-3 loss of function. Strong GCK-1/CCM-3 depletion increased embryonic lethality, with a very low incidence of cytokinesis failure (less than 0.5% of cells, Figure 3C). However, we could not separate this cytokinesis failure from defects in embryo size, since we observe an increase in the incidence of cytokinesis failure in embryos whose size is reduced after GCK-1/CCM-3 depletion suggesting that cytokinesis failure could be secondary to abnormal embryo size. Cytokinetic furrow regression was never observed in zygotes, even in embryos that were significantly smaller than controls. We also never observed polarity defects following 80-90% GCK-1/CCM-3 depletion. Instead, our findings suggest that GCK-1/CCM-3 are downstream of PAR protein-based polarity, and that the primary phenotype resulting from partial loss of GCK-1/CCM-3 during cytokinesis is an increase in equatorial contractility that translates into faster cytokinetic ring constriction. While in the cells we studied, faster cytokinetic furrowing does not translate to cytokinesis failure, this type of dysregulation could be detrimental in other contexts, such as in cells with long chromosomes with point centromeres or other specialized cell types. More generally, CCM proteins including CCM-3 have been implicated in contributing to endothelial integrity by promoting formation and maintenance of tight and adherens junctions (Fischer et al., 2013; Zheng et al., 2010). Future work will be aimed at investigating the intriguing possibility that CCM-3 contributes to endothelial integrity by coordinating junctional dynamics and contractility in the cytokinetic furrow.

## Materials and Methods

### *C. elegans* strains and culture

Worm strains (Table 1) were maintained using standard procedures at 20°C (Brenner, 1974). CRISPR knock-in of GFP at the N-terminus of LET-502 was carried out using an asymmetric repair template consisting of 0.5 kb 3’ to the start codon of *let-502*, and 1.5 kb of the *let-502* coding region. The latter portion of this repair template was obtained from an existing plasmid containing mlc-4p::GFP::LET-502 (pML1595) via removing the C-terminus of *let-502* using Fast Cloning (Li et al., 2011) leaving only 1.5 kb downstream of the start codon. Then 0.5 kb upstream of *let-502* start codon was amplified from genomic DNA (primers: CCTAATCGTTGTCTTTTGATCGGCA and GGCTGCAGCTCGATTTTCGT) to replace the mlc-4p using overlap extension PCR (Bryksin and Matsumura, 2010). To target Cas9 to the *let-502* locus, the sequence of the sgRNA was CGCAGCTCATCCTGCTCCA, which was cloned into pDD162 (Dickinson et al., 2013) and the plasmid (containing Cas9) was injected at concentration of 50 ng/μl. The asymmetric repair template was injected at a concentration of 20 ng/μl, and genome editing was detected via single worm PCR (primers: GCTTGCCTGTCTTATTCATGC (genomic DNA) and TCCGTATGTTGCATCACCTTCACC (GFP)).

**Table 1:**
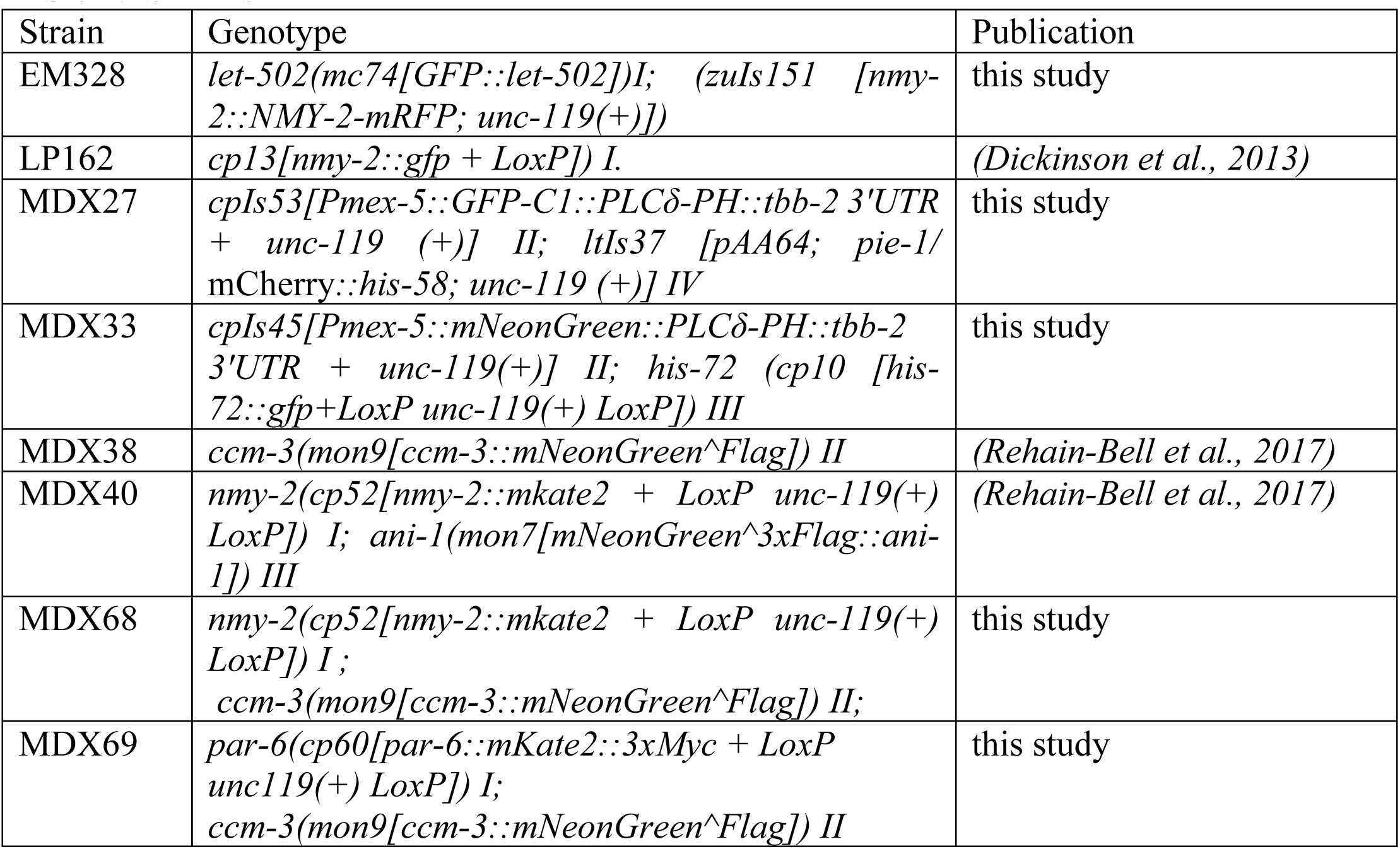
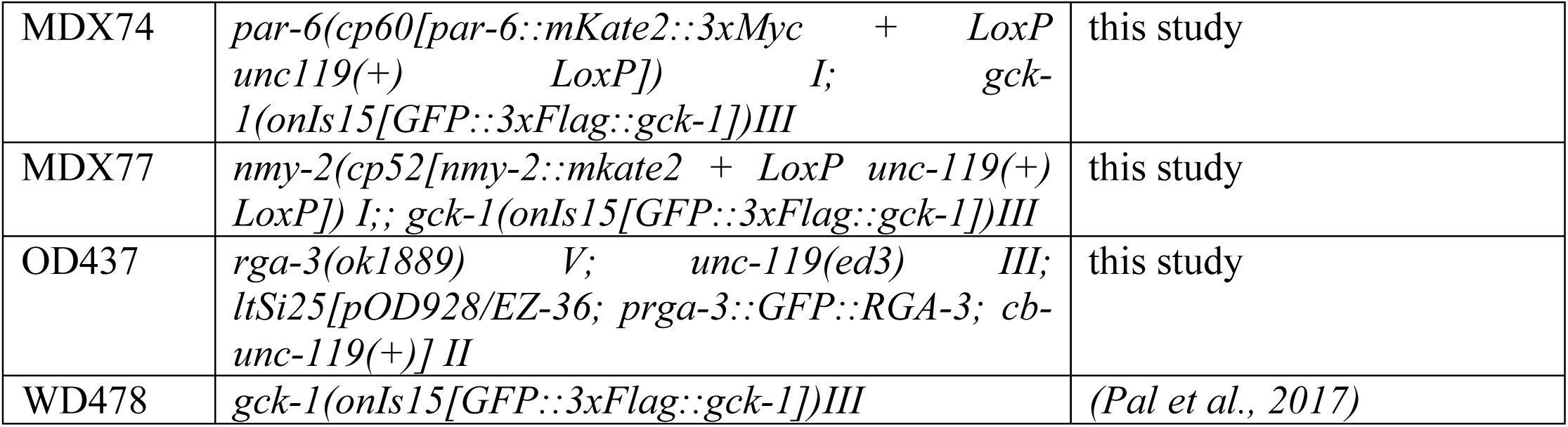
Strains

| Strain | Genotype | Publication |
| --- | --- | --- |
| EM328 | *let-502(mc74[GFP::let-502])I; (zuls151 [nmy-2::NMY-2-mRFP; unc-119(+)])* | this study |
| LP162 | *cp13[nmy-2::gfp + LoxP] I.* | (Dickinson et al., 2013) |
| MDX27 | *cpIs53[Pmex-5::GFP-C1::PLCδ-PH::tbb-2 3'UTR + unc-119 (+)] II; ltIs37 [pAA64; pie-1/mCherry::his-58; unc-119 (+)] IV* | this study |
| MDX33 | *cpIs45[Pmex-5::mNeonGreen::PLCδ-PH::tbb-2 3'UTR + unc-119(+)] II; his-72 (cp10 [his-72::gfp+LoxP unc-119(+)] LoxP) III* | this study |
| MDX38 | *ccm-3(mon9[ccm-3::mNeonGreen^Flag]) II* | (Rehain-Bell et al., 2017) |
| MDX40 | *nmy-2(cp52[nmy-2::mkate2 + LoxP unc-119(+)] LoxP) I; ani-1(mon7[mNeonGreen^3xFlag::ani-1]) III* | (Rehain-Bell et al., 2017) |
| MDX68 | *nmy-2(cp52[nmy-2::mkate2 + LoxP unc-119(+)] LoxP) I;*<br>*ccm-3(mon9[ccm-3::mNeonGreen^Flag]) II;* | this study |
| MDX69 | *par-6(cp60[par-6::mKate2::3xMyc + LoxP unc119(+)] LoxP) I;*<br>*ccm-3(mon9[ccm-3::mNeonGreen^Flag]) II* | this study |
| MDX74 | *par-6(cp60[par-6::mKate2::3xMyc + LoxP unc119(+)* *LoxP]) I; gck-1(onIs15[GFP::3xFlag::gck-1])III* | this study |
| MDX77 | *nmy-2(cp52[nmy-2::mkate2 + LoxP unc-119(+)* *LoxP]) I;; gck-1(onIs15[GFP::3xFlag::gck-1])III* | this study |
| OD437 | *rga-3(ok1889) V; unc-119(ed3) III; ltSi25[pOD928/EZ-36; prga-3::GFP::RGA-3; cb-unc-119(+)] II* | this study |
| WD478 | *gck-1(onIs15[GFP::3xFlag::gck-1])III* | (Pal et al., 2017) |

MDX27 was generated by crossing LP306 (Heppert et al., 2016) to JCC719 (gift from Julie Canman) to introduce mCherry::his58. MDX33 was generated by crossing LP148 (Dickinson et al., 2013) and LP274 (Heppert et al., 2016), MDX68 was generated by crossing MDX38 (Rehain-Bell et al., 2017) and LP229 (Dickinson et al., 2017). MDX69 was generated by crossing MDX38 and LP244 (Dickinson et al., 2017). MDX74 was generated by crossing WD478 and LP244. MDX77 was generated by crossing WD478 and LP229. OD437 was kindly provided by Karen Oegema.

### RNA-mediated interference

Depletions were conducted by feeding worms bacteria, from the Ahringer collection, expressing double-strand RNA (dsRNA) as described previously for 20-24 hours in all conditions except for Figure 3A-C and Supplemental Figure 2A and B for which feeding was carried out for 48 hours (Kamath and Ahringer, 2003; Kamath et al., 2003), For weak *rga-3(RNAi)* experiments, the *rga-3(RNAi)* feeding culture was diluted 5-fold using L4440 and worms were incubated on the resulting feeding plates for 24 hours.

### Quantification of embryonic lethality and frequency of cytokinesis failure

Embryonic lethality was determined by incubating L4 worms on RNAi feeding plates for 24 hours and transferring them to a fresh RNAi plate before incubating for another 24 hours. Parental worms were picked of the plate and embryonic lethality was determined by quantifying the ratio of unhatched embryos to total amount of worms on the plate. For determining cytokinesis failure embryos from MDX27 worms were extrude onto and agarose pad and visually inspected for the presence of multinucleated cells based on DNA morphology. Embryo length was determined in ImageJ measuring the distance from anterior to posterior pole.

### *C. elegans* embryo sample prep and live imaging conditions

Embryos were dissected from gravid hermaphrodites and mounted on 2% agar pads except for Figures 3D-I for which embryos were mounted as inverted hanging drops. Images for Figures 1, 4C-F and Supplemental Figure 4 were acquired with a CoolSNAP Hq camera (Photometrics) mounted on a DeltaVision Image Restoration System (GE) with a 60x/1.42 NA Plan Apochromat objective. For Figure 1 and Supplemental Figure Supplemental Figure B-D, we acquired z-stacks with 1μm spacing through the entire embryo every 20 seconds and every 15 seconds for Figure 4C-E. Data for Supplemental Figure 4A and E-G’ were gathered by taking single images at the midplane of the embryo every 60 seconds.

All other data were acquired on a Nikon A1R microscope body with a 60X 1.27 NA Nikon Water Immersion Objective (Figure 3 B-I and Supplemental Figure 3A and B) or 60x 1.41 NA Nikon Oil Immersion Objective (all other figures) with a GaASP PMT detector using NIS-elements. For Figure 3 D-I, 40 confocal sections with 1 μm spacing through the entire embryo were acquired every 2.7 seconds using the resonance scanner. All other data acquired on the A1R were from single Z-sections at the embryonic cortex with a sampling frequency of 1 second using the Galvano point scanner.

### Segmentation and analysis of fluorescence in the cytokinetic ring

For Figures 1A-B and 3F the ImageJ based software Fiji was used to analyze z-stacks of the full embryo (Rueden et al., 2017; Schindelin et al., 2012; Schneider et al., 2012). First, the cytokinetic ring was isolated. Then, an end-on reconstruction was sum-intensity projected into 2D. The signal in this end-on reconstruction was automatically segmented utilizing the Weka-segmentation plugin. In MATLAB, an ellipsoid was fitted to the segmented signal at each time point giving a measurement of ring perimeter. Fluorescence intensity on the ring at each time-point was measured from sum projections of cytoplasmic-background corrected images where the signal was identified by Weka-segmentation.

For all other quantifications of equatorial fluorescence levels single cortical planes (Figures 2C-E, 5B, C and K and Supplemental Figure 1 C-E) or sum intensity projection of z-stacks (Figure 3H and I) were analyzed in Fiji by preforming a line scan with a width of 5 μm centered around the ingressing furrow. Mean intensity values were calculated and background adjusted using custom MATLAB scripts by subtracting the mean fluorescence intensity of a 5 μm square region in the posterior half of the embryo from the mean intensity data at each time point.

### Analysis of cytoplasmic protein levels and cytokinetic ring enrichment and RNAi depletion levels

Average fluorescence intensity in the cytoplasm was measured using Fiji by measuring the mean fluorescence intensity of a 5 μm by 5 μm square region in center of the embryo. Enrichment on the ring at ∼50% closure was calculated by dividing ring average fluorescence by cytoplasmic average fluorescence at the same time point (Figure 1C-D’). Depletion levels were calculated by preforming a line scan in the equatorial cortex with a width of 5 μm centered around the ingressing furrow and background adjusting by subtracting the mean fluorescence intensity of a 5 μm by 5 μm square region in the cytoplasm

### Line scan analysis and calculation of anterior enrichment

Line scan analysis was performed using Fiji. Lines were drawn along the cortex from the anterior to posterior end of the embryo. The Graphpad software PRISM was then used to fit a sigmoidal line to the data and to calculate the LogIC50. The LogIC50 marks the inflection point where anterior enrichment ends. Anterior enrichment was calculated by determining the ratio of average fluorescence intensity of the most anterior 25% of the cortex to the average fluorescence intensity of the most posterior 25% of the cortex (Supplemental Figure 4).

### Analysis of pseudocleavage furrow depth and persistence

In the MDX33 or MDX27 strains, which express both a membrane and DNA marker, we used Fiji to measure the distance between the two ends of the pseudocleavage furrow at the point of maximal ingression. This number was then subtracted from the total embryo width to determine the depth of the pseudocleavage furrow. To normalize between conditions, we converted the pseudocleavage depth to a percentage of embryo width (Figure 4E). To measure the persistence of the pseudocleavage furrow the time from full regression of the pseudocleavage furrow to pronuclear meeting was calculated (Figure 4D).

### Analysis of total ingression time

We imaged cytokinesis in the strain MDX33 (Figure 3E) or MDX27 (Figure 6A), which expresses both a membrane and DNA marker, with 2.7 second time resolution and calculated the time from initiation defined as the first indentation of the membrane following anaphase onset, to ∼100% closure of the cytokinetic ring, defined as the point when the cytokinetic ring did not decrease in size in the next two time points.

### Tracking and analyzing cortical foci

Kymographs were obtained by performing 2 μm thick line scans along the entire A-P axis of the embryo. Cortical foci during polarization were identified and tracked using Fiji plugin Trackmate (Tinevez et al., 2017). Prior to analysis using Trackmate, single cortical plane images acquired every second were normalized by processing to “enhance contrast” in Fiji. For multi-channel image sequences fluorescence intensities from both channels were summed for each time point to create a binary mask. The normalized and summed stack was combined with the raw data stacks into a hyperstack that was used in Trackmate. Foci were tracked using the normalized and summed stack and values within the tracked areas on the raw data were used for quantitative analysis. We used the Trackmate LoG detector with an estimated foci diameter of 3 μm and median filter thresholding applied in Trackmate to detect foci and the LAP tracer function with maximum linking distance of 2 μm, and “no gap closing distance” to track foci. Processing and analysis of foci properties determined using Trackmate was performed using custom scripts in MATLAB. Mean intensity values within the tracked region were used to determine foci intensity. Mean intensity values were background adjusted at each time point. Background values were obtained by measuring the mean intensity of a 5×5 μm square area in the posterior of the embryo that was devoid of foci for each time point. Only foci persisting for more than 20 seconds were considered in the analysis. Background adjusted intensity values for each track were smoothed using a gaussian filter and aligned at the maximum intensity peak. Mean intensity curves were obtained by averaging intensity values of all aligned foci for a given condition. To compare localization timing to foci, Background adjusted mean intensity values were processed individually for each channel for each track and the time delay for each track was calculated based on the relative time difference the maximum mean intensity within the track was reached.

### Figures and statistical analysis

Figures were generated using Microsoft Excel, MATLAB or GraphPad PRISM software, Statistical significance was determine using a 2-tailed Student’s t-test. Assumptions for normality and equal variance were met for all data analyzed. A p-value of less than 0.05 from a two-tailed t-test was considered significant. Results of the statistical analysis are shown in all figures. All error bars represent standard error unless stated otherwise in the figure legends. Sample size (n) and p-values are given on each figure panel or in the figure legends.

---

**Supplemental Figure 1 (related to Figure 2).**
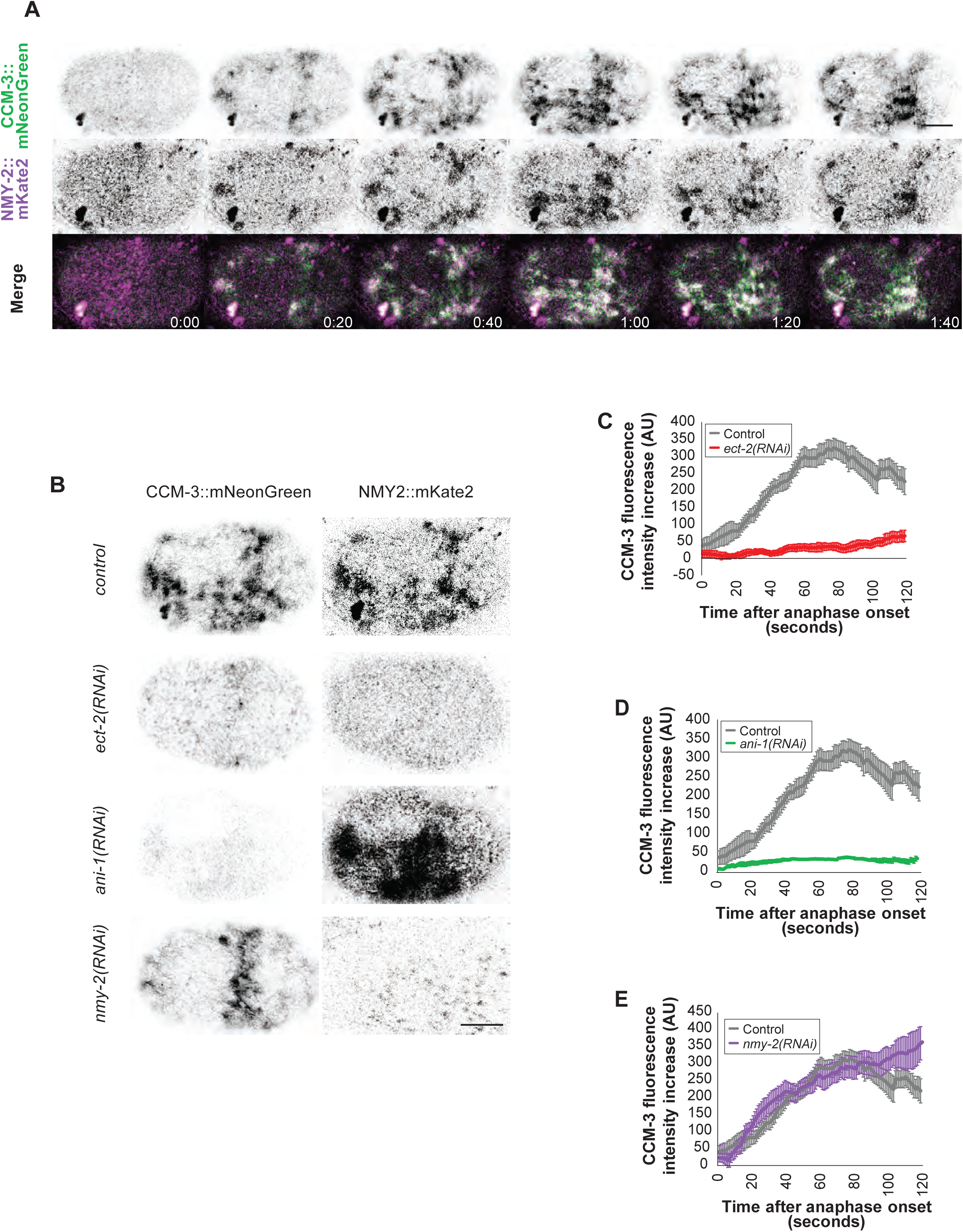
CCM-3 localization is dependent on active RHO-1 and ANI-1 but not NMY-2. A) Representative images of CCM-3::mNeonGreen and NMY-2::mKate localization during anaphase. B) Representative images of embryos expressing both CCM-3::mNeonGreen and NMY-2::mKate in control, *ect-2(RNAi), ani-1(RNAi)* and *nmy-2(RNAi)* embryos 60 seconds after anaphase onset. C-E) Background-adjusted equatorial CCM-3::mNeonGreen levels normalized to that at anaphase onset in control, *ect-2(RNAi)* (C), *ani-1(RNAi)* (D) and *nmy-2(RNAi)* (E) embryos (n=7 for all conditions). (F) Cytoplasmic levels of CCM-3::mNeonGreen in control and ANI-1 depleted cells. Error bars are SE in panels C, D and E, and SD in F.

**Supplemental Figure 2 (related to Figure 3).**
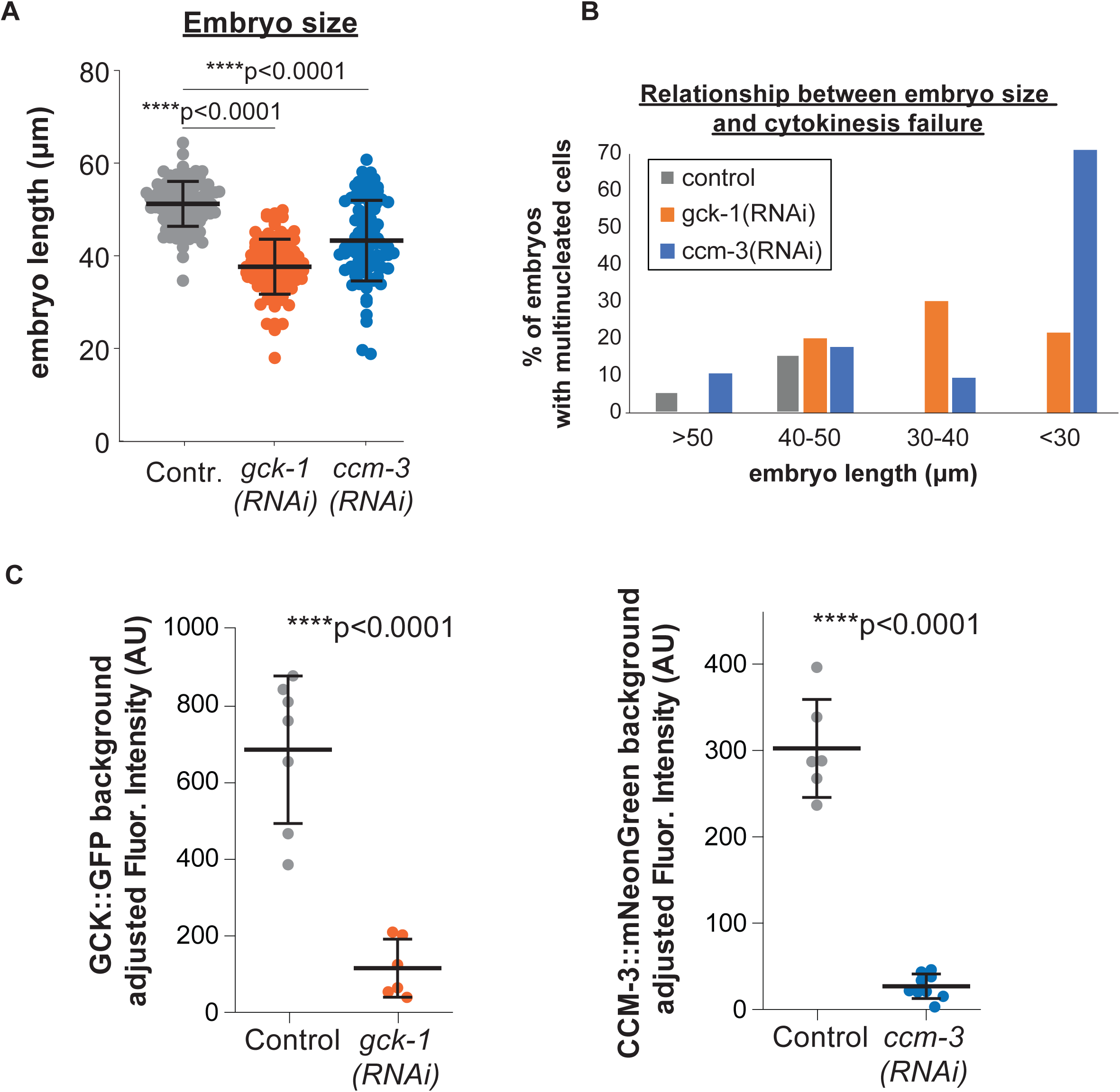
Relationship of embryo size defects and cytokinesis failure in GCK-1 or CCM-3 depleted embryos. A) Quantification of embryo length in control, *gck-1(RNAi)* or *ccm-3(RNAi)* embryos. B) Percentage of embryos with at least one multinucleated cell as a function of embryo length in control, *gck-1(RNAi)* or *ccm-3(RNAi)* embryos. C) Quantification of RNAi depletion levels by comparing fluorescence equatorial fluorescence intensity adjusted for cytoplasmic background in control, *gck-1(RNAi)* or *ccm-3(RNAi)* embryos. Error bars are SD.

**Supplemental Figure 3 (related to Figure 4).**
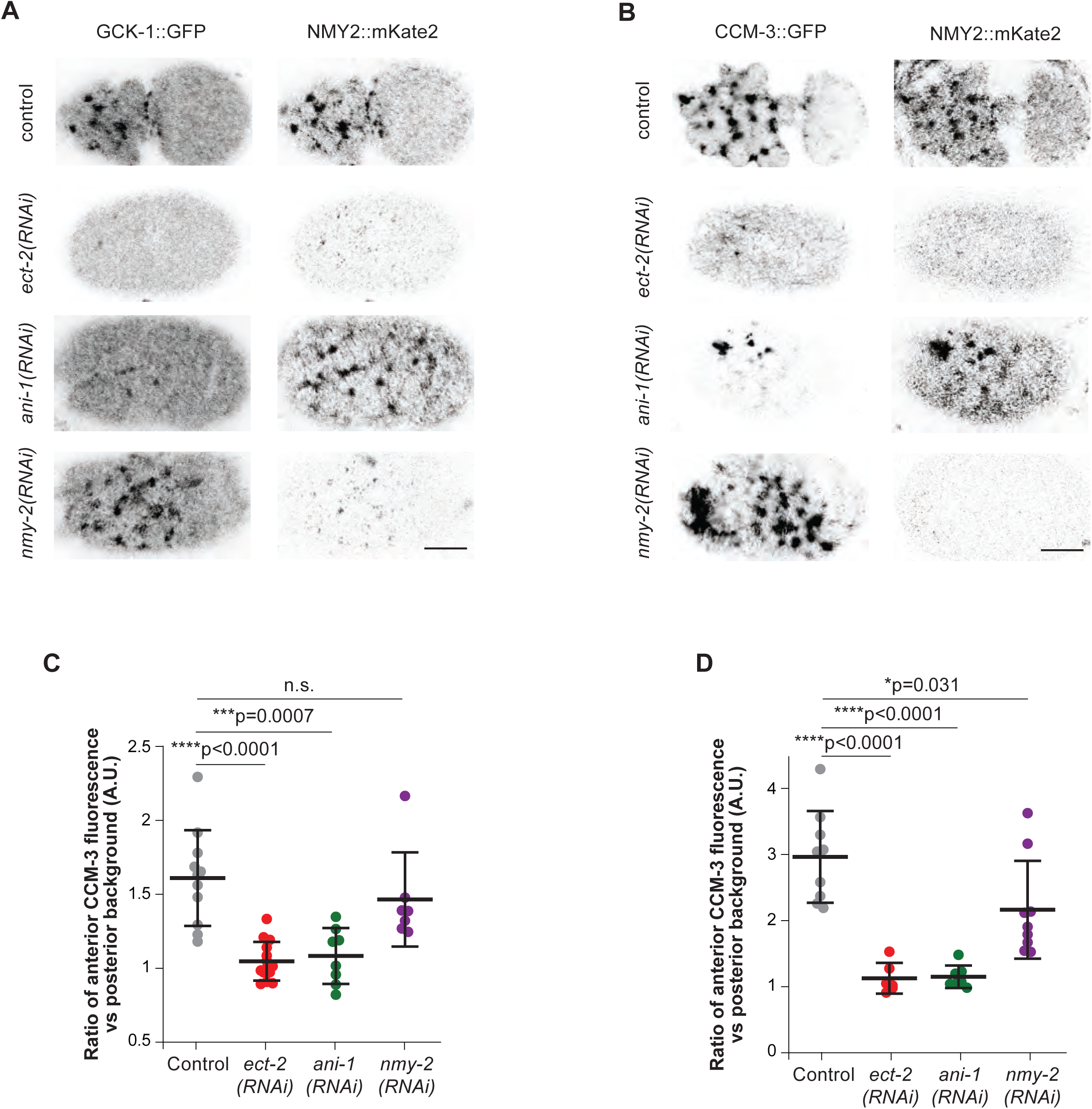
GCK-1 and CCM-3 localization to cortical pulses during polarization are dependent on active RHO-1 and ANI-1 but not NMY-2. Representative images of embryos expressing both GCK-1::GFP and NMY-2::mKate (A) or CCM-3::mNeonGreen and NMY-2::mKate (B) in control, *ect-2(RNAi), ani-1(RNAi)* and *nmy-2(RNAi)* during polarization. Quantification of GCK-1::GFP and CCM-3::mNeonGrean levels in the anterior cortex adjusted for posterior cortical background during polarization in control, *ect-2(RNAi), ani-1(RNAi)* and *nmy-2(RNAi)* embryos. Error bars are SD.

**Supplemental Figure 4 (related to Figure 4).**
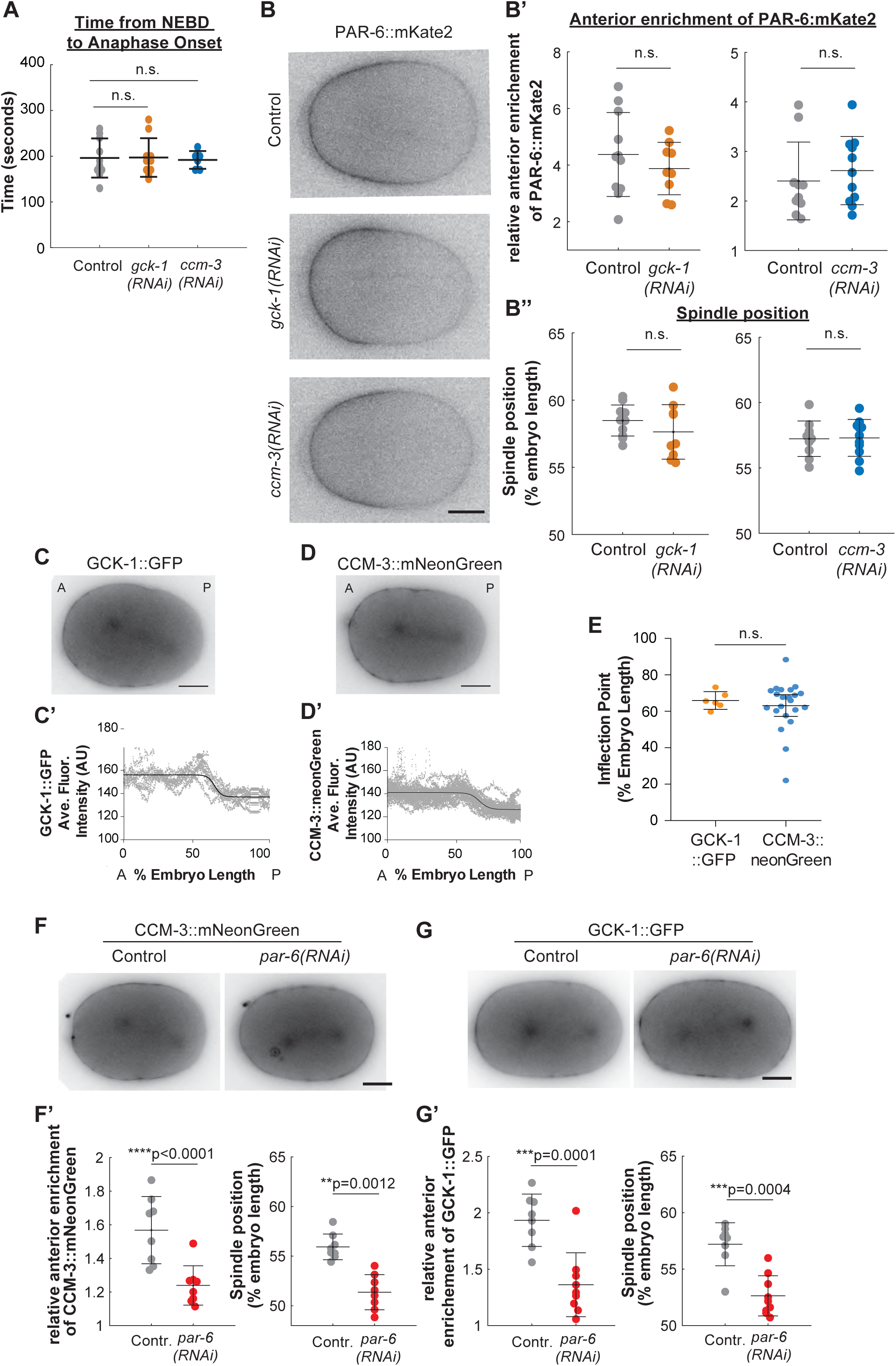
GCK-1/CCM-3 are downstream of polarity. A) Quantification of the time from onset of nuclear envelope breakdown (NEBD) to anaphase in embryos expressing HIS::GFP and mNeonGreen::PH under control, *gck-1(RNAi)* or *ccm-3(RNAi)* conditions. B) Representative single midplane images of PAR-6::mKate2 in control, *gck-1(RNAi)*, and *ccm-3(RNAi)* embryos. B’, B”) Quantification of the effect of *gck-1(RNAi)* and *ccm-3(RNAi)* on anterior enrichment of PAR-6::mKate2 cortical fluorescence (B’) and spindle position (B”). n.s. = no significant difference. C and D) Representative midplane image of GCK-1::GFP localization (C) and CCM-3::mNeonGreen (D) at cytokinesis initiation. A: embryo anterior; P: posterior. C’ and D’) Representative line scans of cortical GCK-1::GFP (C’) and CCM3::mNeonGreen (D’) average fluorescence from the anterior (A) to posterior (P) and sigmoidal fit (LogIC50 = 63.43 and LogIC50 = 61.98 respectively). E) Inflection point (end of the anterior enrichment region; LogIC50) of sigmoidal lines fit to line scan analysis of cortical GCK-1::GFP (n=6 embryos) and CCM-3::mNeonGreen (n=21 embryos) from anterior to posterior. F) Representative single midplane images of CCM-3::GFP in control and *par-6(RNAi)* embryos. F’) Quantification of anterior enrichment of CCM-3::GFP cortical fluorescence and spindle position in control and *par-6(RNAi)* conditions. G) Representative single midplane images of GCK-1::GFP in control and *par-6(RNAi)* embryos. G’) Quantification of anterior enrichment of GCK-1::GFP cortical fluorescence and spindle position in control and *par-6(RNAi)* embryos.

**Supplemental Figure 5 (related to Figure 5).**
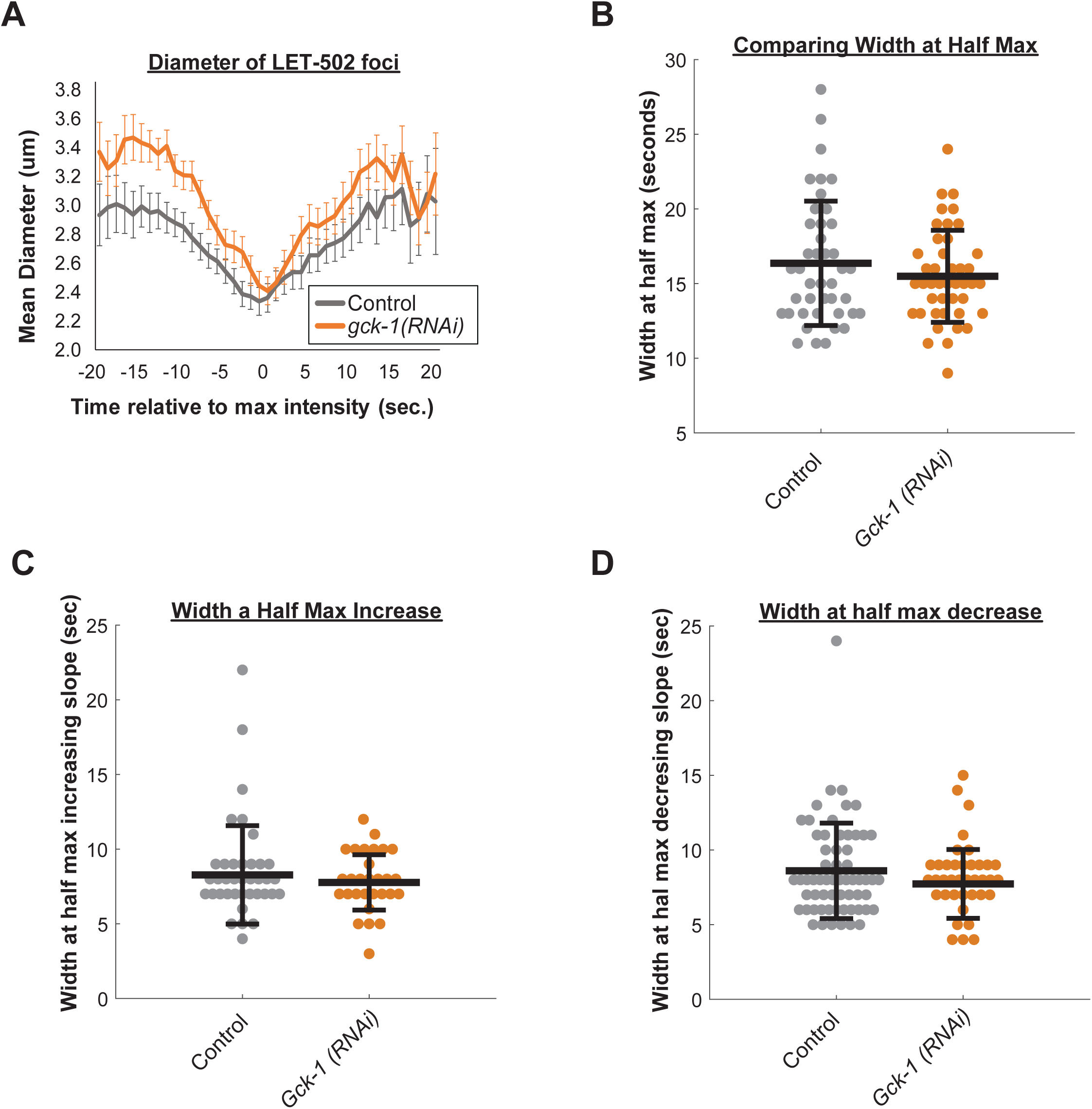
GCK-1 depletion does not change focus dynamics during pulsed contractility. A) Mean estimated diameter profiles of cortical patches based on GFP::LET-502 fluorescence. (B) Width at half max of for all intensity measurements shown in Fig 5E of RHO-1 activity in individual foci in control and *gck-1(RNAi)* embryos. Width at half max measurements of RHO-1 activity increases (C) and decreases (D) in individual patches for control and *gck-1(RNAi)* embryos. Error bars are SD.

**Supplemental movie 1**. Single cortical plane imaging of control embryos expressing GCK-1::GFP (left, green) and NMY-2::mKate (middle, magenta) during anaphase. Acquisition rate 1 frame per second for all Supplemental movies.

**Supplemental movie 2**. Single cortical plane imaging of embryos expressing GCK-1::GFP (left, green) and NMY-2::mKate (middle, magenta) during anaphase after ECT-2 depletion by RNAi.

**Supplemental movie 3**. Single cortical plane imaging of embryos expressing GCK-1::GFP (left, green) and NMY-2::mKate (middle, magenta) during anaphase after ANI-1 depletion by RNAi.

**Supplemental movie 4**. Single cortical plane imaging of embryos expressing GCK-1::GFP (left, green) and NMY-2::mKate (middle, magenta) during anaphase after NMY-2 depletion by RNAi.

**Supplemental movie 5**. Single cortical plane imaging of control embryos expressing GCK-1::GFP (left, green) and NMY2::mKate (middle, magenta) during polarization.

**Supplemental movie 6**. Single cortical plane imaging of control embryos expressing GFP::LET-502.

**Supplemental movie 7**. Single cortical plane imaging of embryos expressing GFP::LET-502 after ECT-2 depletion by RNAi.

**Supplemental movie 8**. Single cortical plane imaging of embryos expressing GFP::LET-502 after RGA-3/4 depletion by RNAi.

**Supplemental movie 9**. Overlay of single cortical plane imaging of control embryos expressing GFP::LET502 (green) and NMY2::mRFP (magenta) with tracked areas identified by ImageJ Trackmate (orange circles) during polarization.

**Supplemental movie 10**. Overlay of single cortical plane imaging of control embryos expressing GCK-1::GFP (green) and NMY2::mKate (magenta) with tracked areas identified by ImageJ Trackmate (orange circles) during polarization.

## Acknowledgements

We thank Julie Canman, Brent Derry, Dan Dickinson, Jenny Heppert, Edwin Munro and Karen Oegema for providing worm strains. We thank Stephan Grill and members of the Maddox labs, especially Paul Maddox, Jenna Perry and Vincent Boudreau, for critical reading and discussion of the manuscript. The authors declare no competing financial interests. KRB was supported in part by a grant from the National Institute of General Medical Sciences under award 5T32 GM007092. This work was supported by European Research Council grant #294744 to ML, and GM102390 from the NIH and 1616661 from the NSF to ASM.

## Author Contributions

Kathryn Rehain Bell and Michael E. Werner contributed to conceptualization, data curation, formal analysis, investigation, supervision, visualization and writing of the original draft. Michael E. Werner also contributed methodology and writing (editing). Anusha Doshi contributed to investigation and visualization. Daniel B. Cortes contributed software and writing (review). Adam Sattler contributed investigation and data curation. Thanh Vuong-Brender contributed a resource. Michel Labouesse contributed to funding acquisition and supervision. Amy Shaub Maddox contributed to conceptualization, funding acquisition, supervision, visualization, and writing (editing).

